# The link between antibiotic resistance level and soil physico-chemical properties

**DOI:** 10.1101/2025.03.17.643646

**Authors:** Mateusz Szadziul, Agata Goryluk-Salmonowicz, Magdalena Popowska

## Abstract

Antimicrobial resistance (AMR) is a critical global health concern. While AMR research has primarily focused on medical and veterinary settings, the spread of antibiotic-resistant bacteria (ARB) and antibiotic resistance genes (ARGs) through natural environments, including soil, remains poorly understood. This study investigates the relationship between soil physico-chemical properties and ARG abundance in environments with varying levels of anthropogenic impact.Soil samples were collected from agricultural fields (both manured and non-manured) and forests, analyzed for 24 physico-chemical parameters, and subjected to DNA extraction. High-throughput qPCR was used to quantify 27 clinically relevant ARGs and 5 mobile genetic elements (MGEs) in the samples. Results revealed significant differences in soil properties between arable and forest soils, particularly in water content, humus levels, sand and silt proportions, and mercury concentration (p≤0.05). Arable soils exhibited a significantly higher abundance of ARGs and MGEs (p=0.0247), with certain resistance genes found exclusively in agricultural environments. Correlation analysis identified strong positive associations between MGEs and ARGs, highlighting the role of genetic elements in AMR dissemination. Additionally, soil properties such as aluminum, nitrogen, and magnesium showed positive correlations with ARG and MGE abundance, while sand content and the carbon-to-nitrogen ratio displayed inverse correlations. The results indicate that heavy metal contamination may play a substantial role in AMR spread through co-selection mechanisms. These findings emphasize the influence of environmental factors on AMR dynamics and highlight the need to integrate soil ecology into AMR mitigation strategies within the One Health framework.

## Introduction

Antimicrobial resistance (AMR) is considered one of the largest health-related issues globally (World Health Organization, 2024). AMR, which exists in both clinical and natural environments, is the subject of numerous studies (Hernando-Amado et al., 2019). However, the issue of AMR spread is described mainly from a medical and veterinary perspective. Reducing antibiotic consumption and implementing other strategies are important measures to lower antimicrobial resistance in a global health context, but the evolution and spread of antibiotic-resistant bacteria (ARB) and antibiotic resistance genes (ARGs) into the environment cannot be reversed (Collignon et al., 2018). Therefore, the dissemination of AMR through the environment leads to various health issues and confirms the present-day consensus that tackling AMR requires action aligned with the ‘One Health’ approach—a holistic framework in which human, veterinary, and environmental settings are interconnected (Aarestrup et al., 2021; Goryluk-Salmonowicz and Popowska, 2022).

Multiple environmental dissemination routes for resistant bacteria have been identified as potentially significant for the proliferation of AMR, particularly in the context of intensive agriculture and increasing urbanization (Thanner et al., 2016). Some critical points have been identified, but the lack of knowledge regarding the propagation and amplification of AMR in the environment complicates AMR prevention and mitigation in human and veterinary medicine (Berendonk et al., 2015). Soil biocenosis is among the most biodiverse systems on Earth, and soils play a crucial role in food production. ARB can disseminate into soils and subsequently be transferred back to humans and animals while evolving and acquiring new ARGs from environmental bacteria in the process. The pool of resistance genes prevalent in the environment, known as the ‘resistome,’ poses a threat to human and animal health if transferred to infectious pathogens (Davies and Davies, 2010; Gillings et al., 2017). Therefore, environments influenced by the input of ARB, which also carry genetic mobility determinants (such as plasmids, transposons, or integrons), should not be regarded merely as passive dissemination routes. Additionally, the presence of pesticides, antibiotics, heavy metals, and other toxic substances polluting the environment may promote the acquisition, emergence, and co-selection of resistance mechanisms against these compounds (Pal et al., 2015; Chen et al., 2019; Bazzi et al., 2020). The accumulation of antibiotics in soils depends on their physico-chemical properties. Furthermore, physico-chemical characteristics of the environment have also been identified as potential factors influencing the abundance of ARGs (Song et al., 2016; Zhang et al., 2019).

Currently, the understanding of the fate of ARB and ARGs released into the environment is limited. Both the transfer of ARGs and the balance between retention and outcompetition of ARB in natural communities remain largely unknown. This is due to the vast array of environmental and ecological factors influencing the spread of ARB and ARGs. In this regard, the biodiversity of a given ecosystem is proposed to be pivotal. Indeed, our previous study revealed that in soil environments, higher biological diversity, evenness, and richness were significantly negatively correlated with the relative abundance of over 85% of the ARGs examined. However, a similar effect was not noticeable in river ecosystems due to their more dynamic nature (Klümper et al., 2024).

To determine whether there is a quantitative and qualitative relationship between ARGs and the physico-chemical properties of soil, research samples were collected from cultivated fields (both manured and non-manured) and forest areas. All collected soil samples were analyzed for their physico-chemical properties, including pH value, water content, humus, sand and clay proportions, total nitrogen, carbon-to-nitrogen ratio, and the presence of various chemical elements, including bioavailable elements and heavy metals. In total, 24 different factors were examined. The resistome was characterized by assessing the abundance of 27 clinically relevant ARGs using high-throughput chip-based qPCR. Additionally, the abundance of mobile genetic elements (MGEs) in the samples was evaluated through 5 marker genes associated with AMR. This study reveals that ARG and MGE distribution varies with soil physico-chemical properties and their origin, with higher abundances linked to arable soils and heavy metal concentrations.

## Materials and methods

### Sample collection

The study was conducted on soil samples collected in autumn 2022 from six areas in Poland with varying levels of anthropogenic activity: four agricultural fields (two manured and two non-manured) representing high activity, and two forests representing low activity. At each location, five soil subsamples were collected along two 10 m diagonals laid in an X-pattern. The subsamples were combined in a sterile plastic bag, homogenized, and transferred to the laboratory. In total, approximately 1 kg of soil was gathered per sampling location. Samples were divided into aliquots of 20 g, sieved (2 mm mesh size) and stored at -20 °C. Table S1 presents the locations of the sampling sites.

### Physico-chemical properties analysis

Collected soil samples were analyzed for their physico-chemical properties by an external laboratory (AGES, Austria). In total, 24 different factors were examined for each soil sample (Table S1).

### DNA isolation

Total DNA was extracted using the DNeasy PowerSoil Pro Kit (Qiagen) according to the manufacturer’s instructions. DNA samples were isolated in four replicates of 0.25 g of soil each and subsequently combined. The quality and quantity of the extracted DNA were assessed using a Qubit 4.0 Fluorometer (dsDNA high-sensitivity assay kit, Invitrogen) and a Colibri spectrophotometer (Titertek Berthold).

### High-throughput qPCR

Determination of the relative abundance of target genes in the samples was performed by Resistomap (Finland) using a SmartChip Real-Time PCR cycler (Takara). A total of 27 ARGs and 5 mobile genetic elements (MGEs) were targeted (Table S3) (Stedtfeld et al., 2018). Quantification included the 16S rRNA gene. The qPCR cycling conditions and initial data processing were carried out as described by Wang et al., 2014. A cycle threshold (CT) of 27 was used as the detection limit (Zhu et al., 2013; Wang et al., 2014). Amplicons with nonspecific melting curves and multiple peaks were discarded from the analysis. The relative abundances of the detected genes, expressed as proportions of the 16S rRNA gene, were calculated using the 2−ΔCT method (Schmittgen and Livak, 2008).

### Statistical analysis

Relationships between physico-chemical properties of soils and the relative abundances of ARGs/MGEs were determined using Spearman’s correlation coefficient and visualized with GraphPad Prism 9. Correlation networks were created using Cytoscape 3.10.1. Calculations included pairwise correlations between physico-chemical properties and ARGs/MGEs as well as correlations among ARGs/MGEs themselves. Correlation was considered strong for a coefficient value of |Rs| > 0.8 and significant for a p-value ≤ 0.05. Additionally, a t-test (Welch’s t-test for single factors and a paired t-test for overall interaction) was performed using GraphPad Prism 9 to assess whether a significant difference existed between arable and forest soils in terms of physico-chemical properties and ARG/MGE abundances. A difference was considered statistically significant for a p-value ≤ 0.05.

## Results

### Physico-chemical properties of soils

Physico-chemical properties were determined across all samples of arable and forest soils, including pH value, water content, proportion of humus, sand, and clay, total nitrogen, carbon-to-nitrogen ratio, and the presence of various chemical elements, including bioavailable elements and heavy metals (Table S1). A comparison of these properties between arable and forest soils was performed using a t-test with Welch’s correction.

The analysis revealed a significant difference between arable and forest soils (p ≤ 0.05) in terms of soil water content, humus, sand and silt proportions, and mercury concentration. Several other factors exhibited weaker but noteworthy trends (p ≤ 0.1), including pH value, total nitrogen content, and the concentrations of magnesium, copper, arsenic, nickel, and aluminum. However, no significant differences (p > 0.1) were found for clay content, carbon-to-nitrogen ratio, or the concentrations of phosphorus, potassium, iron, manganese, zinc, boron, lead, chromium, vanadium, and calcium (Table S2). Overall, forest and arable soils exhibited significantly different physico-chemical properties (p = 0.034; R^2^ = 0.1809).

### Diversity and abundance of ARGs and MGEs

The relative abundance of the analyzed antibiotic resistance genes (ARGs) and mobile genetic elements (MGEs) differed between arable and forest soils (p = 0.0247). The average abundance of these genes was lower in forest soils compared to arable soils (total mean difference = -0.05191 copy numbers/16S rRNA copy number; R^2^ = 0.1524).

Two of the analyzed genes, *blaOXA58* (β-lactam resistance) and *mcr3* (collistin resistance), were not detected in any of the soil samples. Additionally, several genes were found exclusively in arable soils: *intl1_1* (class 1 integron-integrase), *Tn5* (transposase), *aph3-VIa* (aminoglycoside resistance), *ermB_1* (MLSB resistance), *ermF_1, qnrA* (quinolone resistance), *qnrS_1, tetM_2* (tetracycline resistance), *tetO_1* and *dfrA1_1* (trimethoprim resistance). In total, 5 MGEs and 25 ARGs were identified in the analyzed soils (Figure 1). Table S1 presents the relative abundances of the identified genes in soil samples.

**Figure 1.**
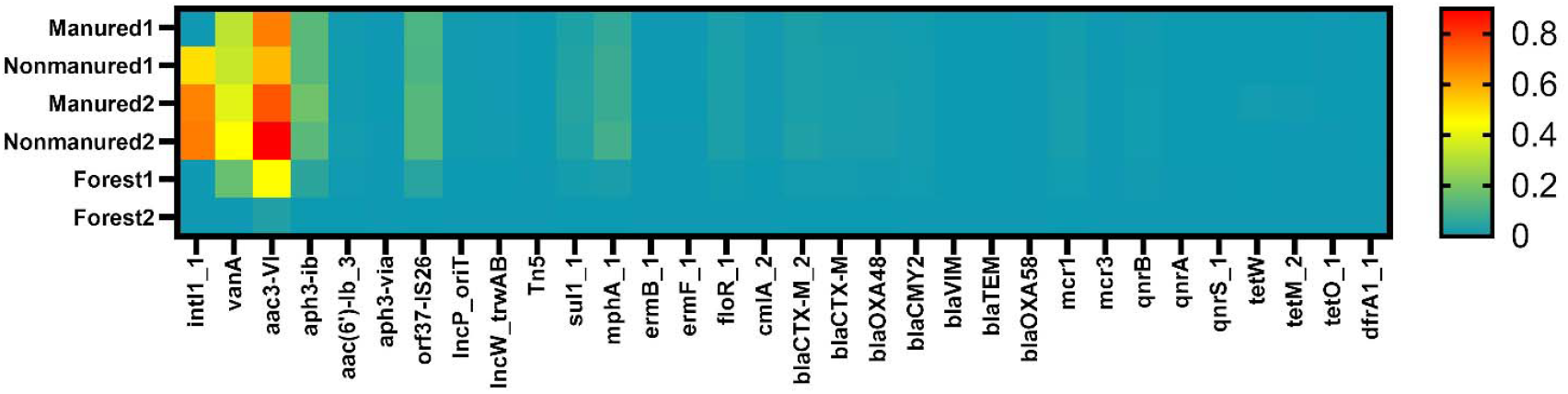
Heatmap of relative ARGs and MGEs abundances in the soils. Values shown are transformed to log10 scale. Samples are color coded from high abundance (red) to below the detection limit (cyan).

A comparison of specific ARG and MGE abundances between arable and forest soils revealed significant differences (p ≤ 0.05) for *sul1_1* (sulfonamide resistance), *mphA_1* (macrolide resistance), *cmlA_2* (chloramphenicol resistance), *blaOXA48* (β-lactam resistance), *blaTEM* (β-lactam resistance), *qnrA, qnrS_1*. Weaker evidence (p ≤ 0.1) suggested a similar difference for *intl1_1, Tn5, aac(6’)-Ib_3* (aminoglycoside resistance), *floR_1* (chloramphenicol resistance) and *blaCTX-M_2* (β-lactam resistance). In all cases, including genes with no significant differences, mean abundances were higher in arable soils (Table S2). However, no significant differences were observed when comparing overall abundances of MGEs or specific antibiotic classes.

### Correlation between ARGs and MGEs

Correlation analysis between mobile genetic elements (MGEs) and antibiotic resistance genes (ARGs) revealed a network consisting of 25 nodes (20 ARGs and 5 MGEs) and 147 edges, representing strong and significant correlations (|Rs| > 0.8; p ≤ 0.05) (Figure 2). All correlations were positive.

**Figure 2.**
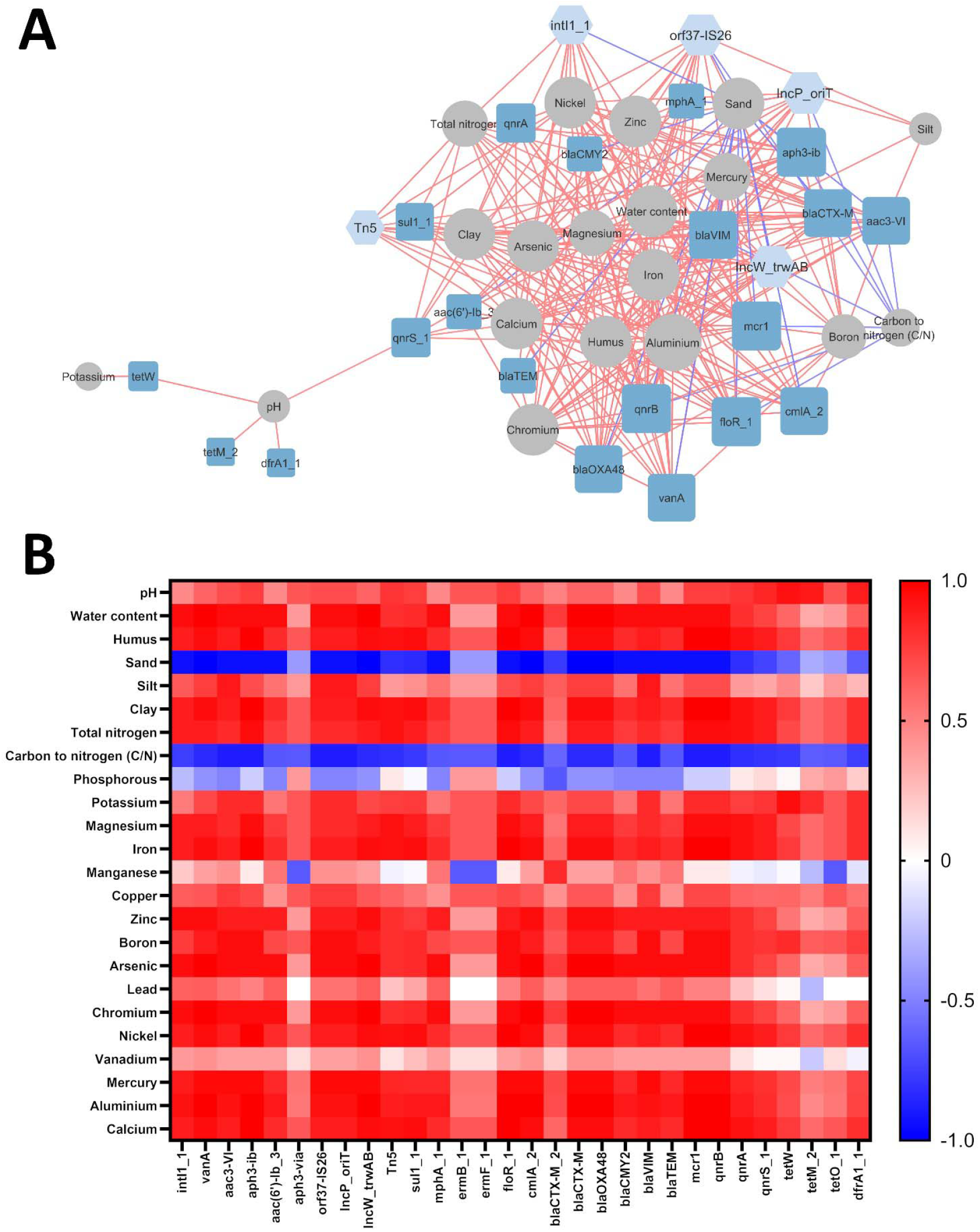
(A) ARG and MGE interaction network in examined soils.The ‘organic layout’ network is presented, where edges indicate strong and significant correlations (|Rs| > 0.8; p-value ≤ 0.05). Positive correlations are shown in red. The size of the nodes represents the degree of interaction, while node colors distinguish MGEs (light blue) from ARGs (dark blue). (B) Heatmap of correlations between ARGs and MGEs, where color represents positive (red) or negative (blue) correlations.

A total of 47 interactions were associated with MGEs, highlighting their role in network formation. All examined MGEs (*intl1_1, orf37-IS26, IncP_oriT, IncW_trwAB* and *Tn5*) interacted with ARGs. However, no significant interactions were observed for the following ARGs identified in soil samples: *aph3-via, ermB_1, ermF_1, blaCTX-M_2* and *tetO_1*.

### Correlation between ARGs/MGEs and physico-chemical properties of soils

A correlation analysis was conducted to examine potential interactions between antibiotic resistance genes (ARGs), mobile genetic elements (MGEs), and various physico-chemical soil properties. The resulting network consisted of 44 nodes (20 ARGs, 5 MGEs, and 19 physico-chemical factors) and 275 edges (excluding interactions between ARGs and MGEs), representing strong and significant correlations (|Rs| > 0.8; p ≤ 0.05) (Figure 3). Of these interactions, 26 were negative, while 249 were positive.

**Figure 3.**
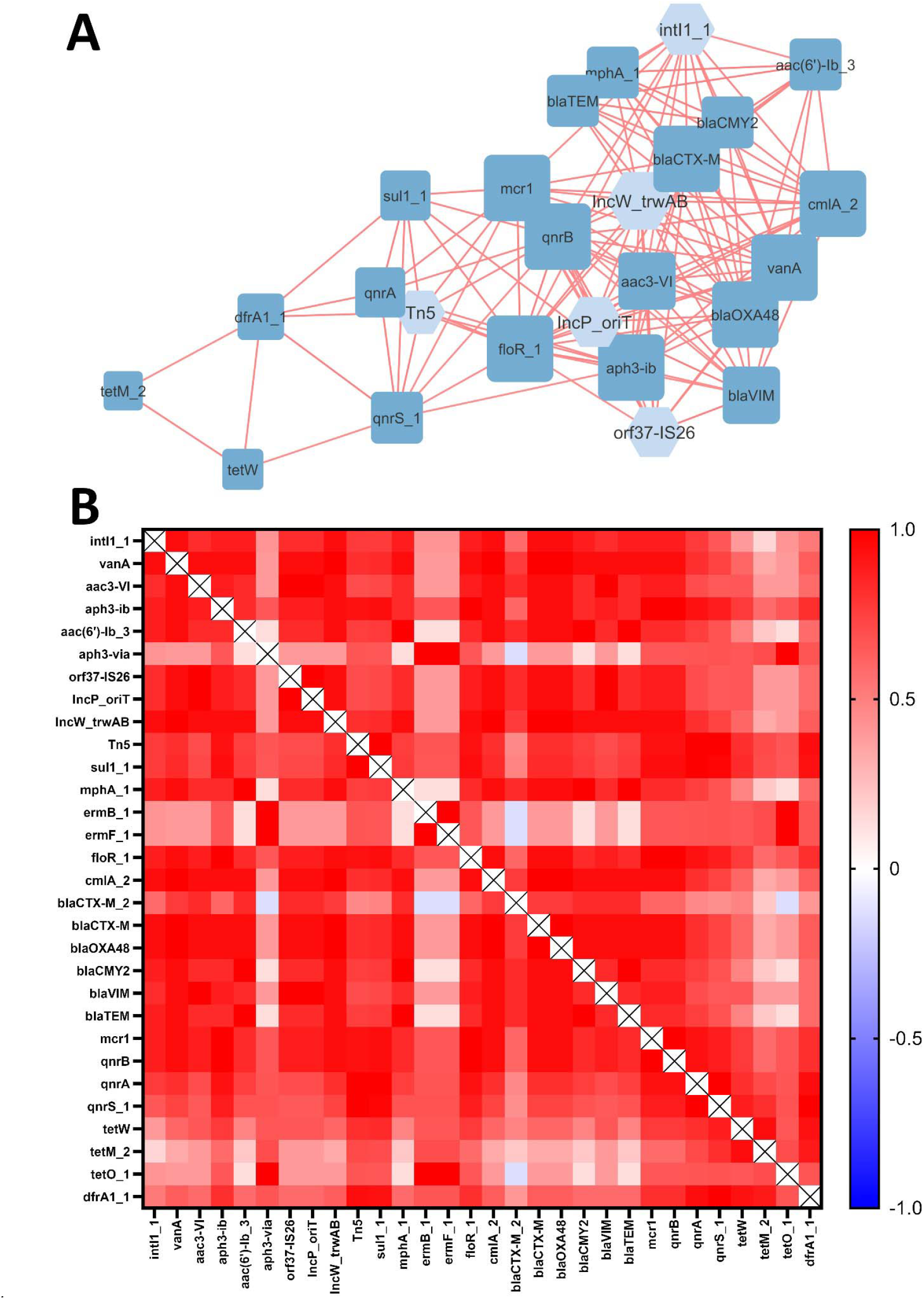
(A) ARG/MGE interaction network with physico-chemical factors in examined soils. The ‘organic layout’ network is presented, with edges indicating strong and significant correlations (|Rs| > 0.8; p ≤ 0.05). Red edges denote positive correlations. Node sizes represent the degree of interaction, while node colors distinguish MGEs (light blue), ARGs (dark blue), and physico-chemical factors (gray). (B) Heatmap of correlations between ARGs/MGEs and soil physico-chemical properties, where red represents positive correlations and blue represents negative correlations.

Specifically, sand content and the carbon-to-nitrogen ratio were inversely correlated with ARG and MGE abundances, whereas most other physico-chemical factors showed positive correlations. No significant correlations were observed between ARG/MGE abundances and copper, lead, manganese, phosphorus, or vanadium. The degree of interaction (number of correlated genes) among physico-chemical factors varied, ranging from 1 to 22 edges. Potassium, silt content, and soil pH exhibited the fewest correlations (1 for potassium and 4 for silt content and pH), while aluminum abundance had the highest number of connections (22).

All examined MGE-encoding genes were correlated with at least one soil parameter. However, five ARGs (*aph3-via, ermB_1, ermF_1, blaCTXM-2* and *tetO_1*) did not exhibit significant interactions with any tested physico-chemical factors.

Aluminum abundance showed a strong positive correlation with all tested MGE-related genes (*intl1_1, orf37-IS26, IncP_oriT, IncW_trwAB, Tn5*). and 17 out of 20 ARGs, excluding *tetW, tetM_2* and *dfrA1_1*. Interestingly, tetracycline resistance genes correlated with only two soil parameters: potassium (*tetW*) and soil pH (*tetW, tetM_2*).

Physico-chemical properties were categorized into three groups based on their correlation profiles with ARGs:

- Group I: Arsenic (As), chromium (Cr), zinc (Zn), and water content. These factors positively correlated with 14 ARGs, including *vanA* (vancomycin resistance), *aac(3)-VI, aac(6’)-Ib_3, aph3-Ib* (aminoglycoside resistance), *mphA* (macrolide resistance), *floR_1, cmlA* (phenicol resistance), *blaCTX-M, blaOXA-48, blaCMY-2, blaVIM, blaTEM* (β-lactam resistance), *mcr1* (colistin resistance), and *qnrB* (quinolone resistance).
- Group II: Calcium (Ca), iron (Fe), nickel (Ni), clay content, and humus content. These factors positively correlated with 13 ARGs, including *vanA, aac3-VI, aph3-Ib, floR_1, cmlA, blaCTX-M, blaOXA-48, blaVIM, mcr1, qnrA, qnrB, qnrS*, and *sul1*.
- Group III: Magnesium (Mg) and nitrogen (N). These parameters had nearly the same correlation profile as Group II, except they lacked interactions with *aac3-VI* and *blaVIM*.

Boron and mercury were also analyzed separately. Boron showed positive correlations similar to those in Group II, except for *sul1_1* and *qnrS*. Mercury, however, was positively correlated with all genes in the boron group as well as *qnrA*.

All MGE-related genes were positively correlated with Group II parameters (clay, humus, Ca, Fe, and Ni). Additionally, four out of five MGEs (*intl1_1, orf37-IS26, IncP_oriT, IncW_trwAB*) correlated with aluminum and Group I parameters (water, As, Cr, Zn). Nitrogen and magnesium correlated positively with *intl1_1, IncW_trwAB*, and *Tn5*, while boron and mercury were associated with *orf37-IS26, IncP_oriT*, and *IncW_trwAB*.

## Discussion

ARGs presence in the soil environment is a complex issue. Various interlinked factors contribute to the distribution of ARGs in soils, including antibiotic residues, anthropogenic pressure, microbial community structure, and soil physico-chemical properties (Čermák et al., 2008; Popowska et al., 2012; Xie et al., 2018; Cycoń et al., 2019). In our study, we analyzed the relationship between ARGs/MGEs presence and soil physico-chemical properties.

The differences in the physico-chemical properties of forest and arable soils were likely influenced by anthropogenic activities in agricultural soil, including fertilization, which may explain the lower total nitrogen levels in forest soils. Previous studies have reported that fertilization affects soil physico-chemical properties (Li et al., 2018; Zhang et al., 2023a). Additionally, agricultural soils may be more sensitive to external inputs, leading to significant changes in their physico-chemical properties compared to forest soils (Bang-Andreasen et al., 2020).

In general, highly fertile soils are typically used for agriculture, whereas forest soils exhibit a broader range of physico-chemical properties compared to arable land. Forest soils generally have lower pH values and fewer available nutrients, which aligns with our findings (Plassart et al., 2019). Our study revealed that sand content was higher (82.3–92.8%) and silt content lower (2.7–12.6%) in forest soils compared to arable soils (34.8–65.9% and 27.7–53.4%, respectively). However, the variation in clay content between the examined soils was minimal, and no significant differences were observed.. Previous research has reported higher silt and clay contents in arable soils compared to forest soils (Tewari et al., 2016). Polish soils generally have high sand content, and its lower levels in arable soils could be attributed to continuous agricultural use, increased fertility, and fertilization practices (Ballabio et al., 2016). Water content was also significantly lower in forest soils in our study. In addition to anthropogenic influences such as field irrigation, the sandy soil structure of forests, characterized by low compaction resistance and macropore formation, may also be a contributing factor (Gregory et al., 2007).

Our previous research on forest soils across several European countries identified the most abundant ARGs as *aac3-VI* (aminoglycoside resistance), *dfrA1* (trimethoprim resistance), *mphA* (MLSB) and *qnrS* (quinolone resistance) (Klümper et al., 2024).In this study, we obtained similar results, but with a few significant differences: *vanA* (vancomycin resistance) was the most abundant ARG, while *dfrA1* was the least abundant ARG. Other studies on ARGs in agricultural soils have shown that genes conferring resistance to ß-lactams, glycopeptides (e.g., vancomycin), and aminoglycosides comprise up to 35% of detected ARG subtypes, which aligns with our findings, although with lower ß-lactamase abundances (Zhang et al., 2021). Our study revealed that ARGs and MGEs are more abundant and diverse in arable soils, suggesting that agricultural areas are important hotspots for ARG pollution (Sun et al., 2020). However, while manure application plays a key role in ARG distribution and increasing abundance in arable fields, we did not observe this relationship in our data (Huang et al., 2021; Zalewska et al., 2021, p.; Zhang et al., 2023b).This may be due to the unknown fertilization history of the examined fields, as manure application from previous years could still impact ARG abundance (Zhang et al., 2023b). Another possible explanation is that extreme levels of metal concentrations contributed to ARG elevation in one set of soils via co-selection mechanisms (Zhang et al., 2018; Goryluk-Salmonowicz and Popowska, 2022; Zou et al., 2025). Similar evidence exists for the co-selection of MGEs in metal-polluted soils (Zhao et al., 2019). Further studies are needed to elucidate the mechanisms of ARG and MGE co-selection in soils affected by significant metal pollution.

Our analysis revealed a substantial number of correlations between ARGs and MGEs, with all MGEs linked to at least one ARG, suggesting their role in establishing genetic networks. Indeed, in many environments crucial for antibiotic resistance spread, MGEs are key contributors to ARG abundance (Yang et al., 2021; Delgado-Baquerizo et al., 2022; Li et al., 2024). According to our findings, the presence of 20 out of 27 examined ARGs was positively correlated with other ARGs. Research on Chinese soils heavily polluted with industrial waste also identified similar coexistence patterns between ARGs and MGEs, with single genes linked to up to 13 other ARGs (Yang et al., 2021). Multiple correlations between ARGs and MGEs were also observed in Northern Ireland, where several ARGs corresponded to class 1 integrons and transposons, with intl1_1 showing the highest number of correlations, in agreement with our results (Zhao et al., 2019). Among the ARGs examined in this study, *vanA* exhibited the highest number of interactions (16) with other ARGs/MGEs. This could be attributed to the widespread presence and relatively high abundance of vancomycin resistance genes in the environment (Han et al., 2022; Zheng et al., 2022; Zhang et al., 2025). Previous study has suggested that high ARG abundance is associated with decreased phylogenetic diversity (Klümper et al., 2024). Future research should further investigate the relationship between ARG and MGE abundance and microbiome composition.

The presence of antibiotic resistance genes may be influenced by soil metal concentrations and other elemental factors. In our study, one of the arable soils exhibited extremely elevated concentrations of zinc, lead, and chromium, alongside the highest ARG/MGE abundances among all examined soils. While the direct effect of soil physico-chemical properties and heavy metal presence on ARG variation may be small compared to the impact of microbial community structure (3.57% and 1.19% versus 79.76%, respectively), these factors significantly shape the bacterial community, making soil properties a major contributor to ARG and MGE profile variations (Yang et al., 2021; Zhang et al., 2021).

In this study, aluminum content emerged as the most significant physico-chemical factor associated with increased ARG and MGE abundances. Several studies have described the role of aluminum compounds in facilitating antibiotic resistance gene transfer under laboratory conditions (Liu et al., 2019; Aras et al., 2023). A soil study by Ferreira et al., 2024 found that aluminum presence was positively correlated with *tetX, sul1* and *mphA* gene abundance. Similarly, an analysis of residential soil samples in Australia identified aluminum as a major factor influencing ARG abundance, with aluminum levels correlating with the presence of *blaOXA, blaTEM, tetM, tetW, sul2* and *sul3* (Knapp et al., 2017). Our study, conducted on soils in Poland, supports these findings and confirms the consistency of this relationship.

Knapp et al., 2011 analyzed Scottish soils and identified correlations between 11 ARGs and soil metals. Two ß-lactam resistance genes (*bla*CTX-M and *bla*OXA48) were positively correlated with chromium content (53.1 mg/kg), aligning with our findings (mean chromium content: 29.8 mg/kg). However, we observed no correlation between tested ARGs and elements such as copper (1–8.8 mg/kg), lead (9–700.4 mg/kg), manganese (3.76–308 mg/kg), phosphorus (61–117 mg/kg), or vanadium (6.14–24.4 mg/kg). These results are consistent with other studies; Knapp et al., 2011 also observed no correlation of ARGs with lead in concentrations of 10 – 1000 mg/kg. also found no correlation between ARGs and lead in concentrations of 10–1000 mg/kg. Despite existing evidence that copper exposure may act as a selective pressure for antibiotic resistance, our study did not find a significant relationship between copper content and ARG/MGE abundance (Kang et al., 2018).

In summary, most physico-chemical properties showed a positive correlation with all examined ARGs and MGEs. This study provides a framework for assessing the relationship between ARGs, MGEs, and soil physico-chemical parameters, which could inform the development of targeted fertilization and remediation strategies to mitigate the environmental impact of AMR. Therefore, further research is needed to thoroughly investigate the influence of specific physico-chemical properties on ARG and MGE abundance. Given that these properties shape the entire soil ecosystem, future studies on ARG pollution should explore the interplay between soil characteristics, microbiome composition, and ARG diversity and abundance.

## Conclusion

This study highlights the differences in ARG and MGE distribution based on soil physico-chemical properties and their origin. Forest and arable soils exhibit distinct physico-chemical characteristics, with ARGs and MGEs generally more abundant in arable soils. Strong and significant correlations between MGEs and ARGs were observed across all soils. The results indicate that soil physico-chemical properties influence ARG and MGE abundances, with nearly all examined properties associated with specific ARGs and MGEs. In most cases, these correlations were consistently positive or negative across all genes, with aluminum abundance showing the highest number of associations, reinforcing the role of metal pollution in promoting AMR spread. A comprehensive assessment of AMR’s impact on soil environments requires further research examining the interaction between all contributing factors, including physico-chemical properties, climatic conditions, microbiome composition, and anthropogenic influences such as fertilization and contamination.

## Author Contributions

M.SZ.—methodology, formal analysis, investigation, data curation, writing—original draft, visualization; A.G-S.—formal analysis, investigation, writing—original draft; M.P.— conceptualization, writing—review and editing, project administration, funding acquisition.

## Data availability statement

The datasets analyzed in this study are included within the article and its supplementary files or, if not, are available from the corresponding author on reasonable request.

